# A Neanderthal/Denisovan GLI3 variant contributes to anatomical variations in mice

**DOI:** 10.1101/2023.07.03.547394

**Authors:** Ako Agata, Satoshi Ohtsuka, Ryota Noji, Hitoshi Gotoh, Katsuhiko Ono, Tadashi Nomura

**Affiliations:** Developmental Neurobiology, Kyoto Prefectural University of Medicine, Japan; Laboratories for Experimental Animals, Kyoto Prefectural University of Medicine, Japan; Applied Biology, Kyoto Institute of Technology, Japan

**Keywords:** GLI3, Neanderthals, Denisovans, skeletal development, evolution

## Abstract

Changes in genomic structures underlie phenotypic diversification in organisms. Amino acid-changing mutations affect pleiotropic functions of proteins, although little is known about how mutated proteins are adapted in existing developmental programs. Here we investigate the biological effects of a variant of the GLI3 transcription factor (GLI3^R1537C^) carried in Neanderthals and Denisovans, which are extinct hominins close to modern humans. R1537C does not compromise protein stability or GLI3 activator-dependent transcriptional activities. In contrast, R1537C affects the regulation of downstream target genes associated with developmental processes. Furthermore, genome-edited mice carrying the Neanderthal/Denisovan GLI3 mutation exhibited various alterations in skeletal morphology. Our data suggest that an extinct hominin-type GLI3 contributes to species-specific anatomical variations, which were tolerated by relaxed constraint in developmental programs during human evolution.

## Introduction

Understanding the genetic mechanisms underlying phenotypic variation is a fundamental challenge in developmental and evolutionary biology. Several empirical studies suggest that adaptive mutations occurring in *cis*-regulatory regions are a major force driving morphological evolution (1–4). In contrast, mutations in protein-coding regions have been thought to play little role in anatomical variations, since structural mutations often disrupt protein function and cause severe malformations or pathological conditions in organisms. However, recent studies have revealed that changes in protein structures result in novel pleiotropic functions (5–7). Thus, nucleotide variants that alter amino acid sequences contribute significantly to phenotypic diversity.

Adaptive mutations in coding regions are behind the phenotypic diversifications in human evolution as well (8–11). Accumulating data have shown that several admixtures resulted in gene flow between modern human ancestors and archaic hominins such as Neanderthals and Denisovans, who diverged from modern humans approximately 520-630 thousand years ago (12, 13). Fossil evidence has indicated distinct anatomical characteristics in Neanderthals compared to modern humans, including elongated and low crania, larger brow ridges, and broader rib cages (14). Putative morphological profiles inferred from DNA methylation patterns predict that Denisovans may have shared some Neanderthal traits (15). Some of the genetic diversity of extant humans is derived from the extinct hominins (12, 16–19). Recent studies have unveiled some of the functional consequences of coding variations specific to archaic or modern humans (20–25). However, the impact of archaic genomic variants in anatomical structures has not been directly demonstrated.

Previous studies have revealed that relative to modern humans, all sequenced Neanderthals and Denisovans carry an amino acid substitution in the C-terminal region of GLI3 (R1537C) (12, 26). GLI3 is a GLI-Krüppel family transcription factor that plays essential roles in embryonic development by mediating Hedgehog signaling (27). In the absence of Hedgehog, GLI3 is cleaved to form the N-terminal repressor form, while the presence of Hedgehog prevents protein processing, which triggers intracellular accumulation of full-length GLI3. Functional disruptions of GLI3 normally result in severe abnormalities in various anatomical structures in mice and humans (28–30). However, the consequences of the amino acid substitution of the GLI3 protein and phenotypic outcomes have not been clarified.

Here, we investigated the biological effects of the Neanderthal/Denisovan GLI3 variant (R1537C). Our study shows that R1537C substitution affects the regulation of downstream target genes associated with developmental processes without interfering Hedgehog-dependent transcriptional activities. Genome-edited mice carrying the Neanderthal/Denisovan GLI3 mutation exhibited various alterations in skeletal morphology depending on genetic backgrounds. The data suggest that the extinct hominin-type GLI3 variant affects species-specific anatomical variations, which were accommodated by concurrent changes in developmental constraints during human evolution.

## Results

### The R1537C variant might be accommodated in archaic and modern humans

R1537C is localized at C-terminal transactivation domain 1 (TA1) of human GLI3 (Figure 1A). At this position, arginine (R) is extensively conserved among vertebrates (Figure 1B), suggesting that the corresponding arginine residue is a derived mutation in Neanderthal/Denisovan lineages. *In silico* analysis of protein functions predicted that R1537C is deleterious to human GLI3 (Figure 1B). Of note, the SIFT program suggested that substitution to histidine was tolerable at this position, which corresponds to the altered amino acid in Koala Gli3 (Figure 1B). These data suggest that relaxed constraint on the residue occurred in the evolution of archaic human lineages. Next, we investigated the R1537C variant exists in modern human population. Present-day human genomes outside Africa contain 1-4% DNA derived from Neanderthals, while approximately 6% of the Melanesian genomes are derived from Denisovans (16, 31). We confirmed that the single nucleotide polymorphism (SNP) corresponding to the R1537C variant also exists in the modern human population [rs35364414 (G/A)]. The allele frequency of the R1537C variant (A allele) ranges from highest in the European population (3.7%-7.7%) to lowest in the African population (0.8%-2.1%) (The 1000G genome dataset, Figure 1C). A previous study suggested that the chromosomal fragment containing the SNP was inferred to be derived from archaic hominins (19). These lines of evidence suggest that the R1537C variant is accommodated in genetic background of archaic and present-day humans.

**Fig. 1.**
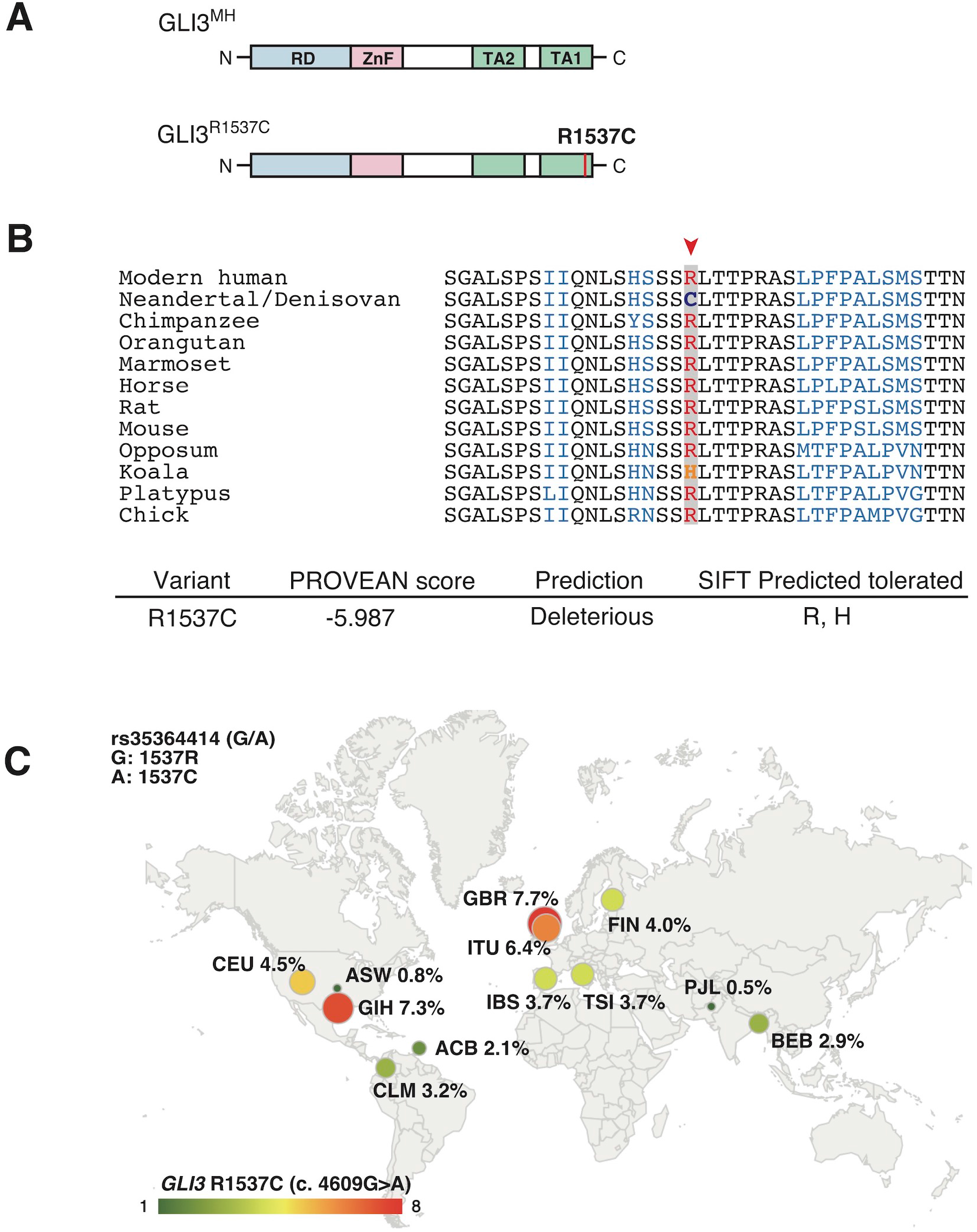
GLI3^R1537C^ is a derived mutation in archaic and present-day humans. (A) Protein structures of the modern human GLI3 (GLI3^MH^) and a Neanderthal and Denisovan GLI3 variant (GLI3^R1537C^). RD: repressor domain; ZnF: Zinc finger type DNA binding domain; TA1: transactivation domain 1; TA2: transactivation domain 2. (B) Upper panel: amino acid sequences of GLI3/Gli3 C-terminal regions in different mammalian species. A red arrowhead indicates the corresponding arginine position. Lower panel: PROVEAN score and SIFT prediction of GLI3^R1537C^. (C) Allele frequencies of R1537C (rs35364414; A allele) in 1000G populations. ACB: African Caribbean in Barbados, ASW: African ancestry in Southwest US, CLM: Colombian in Medellin, Columbia, CEU: Utah residents with Northern and Western ancestry, FIN: Finnish in Finland, GBR: British in England and Scotland, IBS: Iberian population in Spain, TSI: Toscani in Italy, BEB: Bengali in Bangladesh, GIH: Gujarati Indian in Houston, ITU: Indian Telugu in the UK, PJL: Punjabi in Lahore, Pakistan, STU: Sri Lanken Tamil in the UK.

### The R1537C variant does not compromise protein stability and Hedgehog-dependent GLI3 functions

Next, we investigated whether R1537C substitution affects GLI3 protein expression. To test this, we overexpressed modern human-type GLI3 (GLI3^MH^) or a Neanderthal/Denisovan GLI3 (GLI3^R1537C^) in human embryonic kidney cells (HEK293T cells) and performed Western blotting. The amounts of full-length and truncated forms of GLI3 (GLI3 full and GLI3R, respectively) were comparable between GLI3^MH^ and GLI3^R1537C^ (Figure 2A, B). To further examine the stability of GLI3^MH^ and GLI3^R1537C^, we overexpressed luciferase tagged GLI3^MH^ and GLI3^R1537C^ and measured luciferase activity after halting protein expression. We confirmed that the degradation rate was not different between of GLI3^MH^ and GLI3^R1537C^ (Figure 2C). Thus, in contrast to *in silico* prediction, R1537C substitution does not disturb the protein stability of GLI3.

**Fig. 2.**
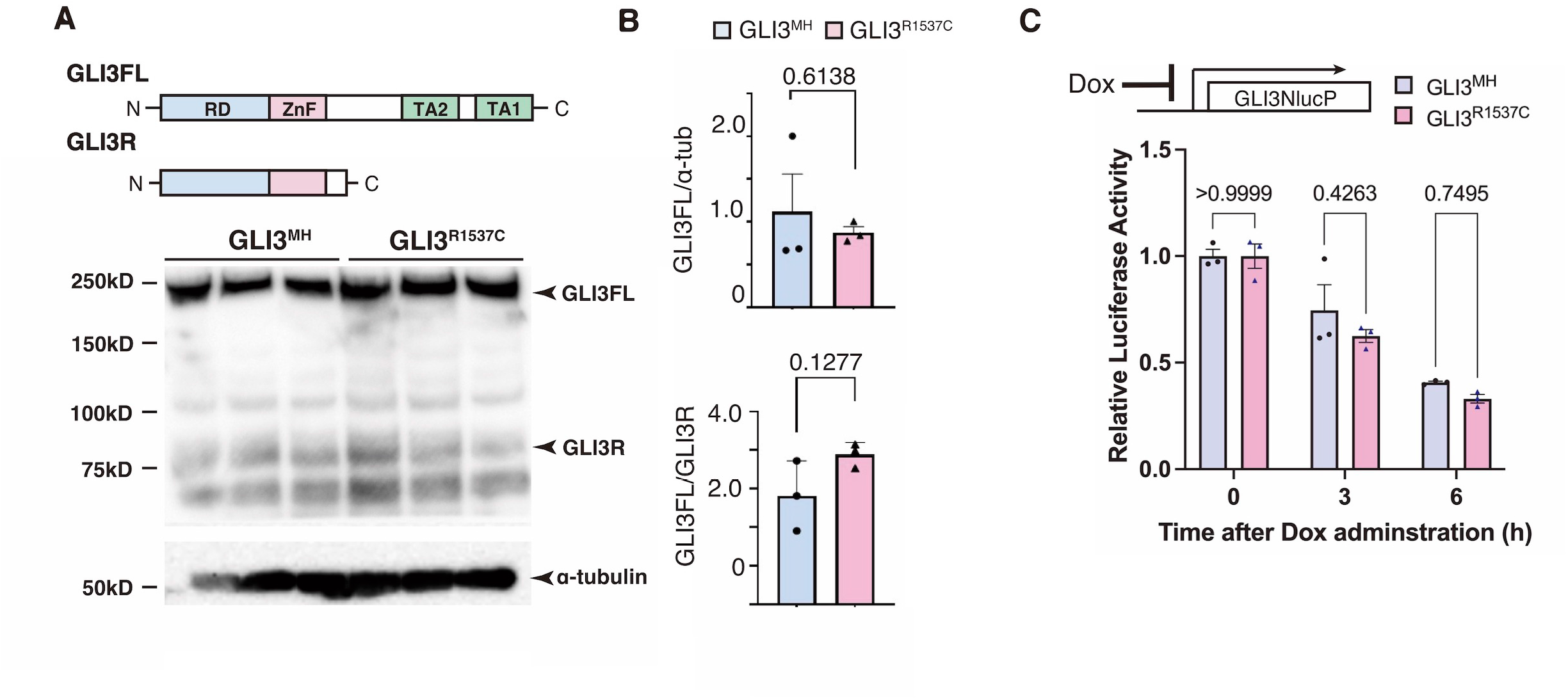
R1537C does not compromise GLI3 protein stability. (A) Western blotting of GLI3^MH^ and GLI3^R1537C^. (B) Quantification of GLI3FL (full length, upper panel) and GLI3FL/GLI3R ratio (lower panel). (C) Comparison of the degradation speed of nanoluc (NlucP)-tagged GLI3^MH^ and GLI3^R1537C^ after halting protein synthesis by doxycycline administration.

To investigate the R1537C affects transcriptional activity of GLI3, we introduced a reporter vector containing GLI-binding sequences together with GLI3^MH^ or GLI3^R1537C^ into HEK293T cells and performed luciferase assay. We found that introduction of GLI3^R1537C^ significantly decreased reporter activity compared to GLI3^MH^ (Figure 3A), suggesting that R1537C alters the transcriptional potential of full-length GLI3.

**Fig. 3.**
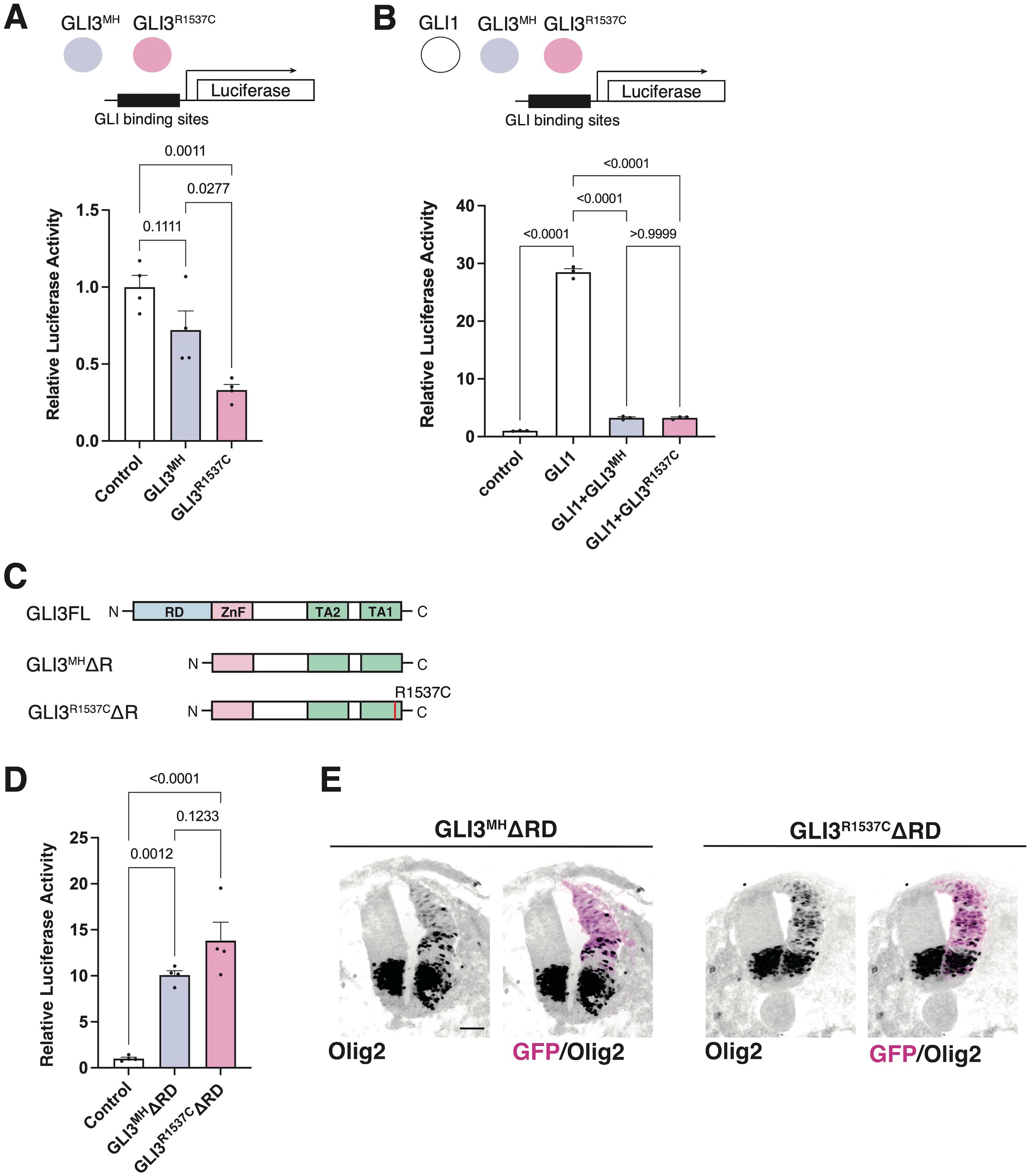
R1537C does not interfere repressor or activator-dependent transcriptional activities. (A) GLI-reporter activities in HEK293T cells overexpressing GLI3^MH^ or GLI3^R1537C^. (B) GLI-reporter activities in HEK293T cells overexpressing GLI3^MH^ or GLI3^R1537C^ together with GLI1. (C) Protein structures of full-length and repressor domain-truncated GLI3 (GLI3^MH^ΔRD or GLI3^R1537C^ΔRD). (D) GLI-reporter activities in HEK293T cells overexpressing GLI3^MH^ΔRD or GLI3^R1537C^ΔRD. (E) Induction of Olig2 expression in the developing chick neural tube by overexpression of GLI3^MH^ΔRD or GLI3^R1537C^ΔRD. Scale bar: 50 µm (E).

Next, we test whether R1537C substitution affect GLI-mediated Hedgehog signaling. Previous studies reported that full-length GLI3 predominantly acts as an inhibitor of Hedgehog signaling, as GLI3 negatively regulates GLI1-dependent transcriptional activity (32). Thus, we examined the effect of R1537C on the suppressive potential of GLI3 in the presence of GLI1. To test this, we overexpressed a GLI reporter vector together with GLI1 and GLI3^MH^ or GLI3^R1537C^ in HEK293T cells (Figure 3B). Compared to the control, GLI1 strongly enhanced reporter activity, while co-transfection of GLI3^MH^ or GLI3^R1537C^ significantly decreased GLI1-dependent reporter activity, indicating that the repressive activity of GLI3 is not curtailed by the amino acid substitution (Figure 3B).

Full-length GLI3 contains both transcriptional repressor and activator domains. To directly assess the effect of R1537C on the transcriptional activator domain, we removed the N-terminal repressor domain and examined luciferase activity in response to GLI3 variants (GLI3^MH^ΔRD and GLI3^R1537C^ΔRD, Figure 3C). Compared to the control, both GLI3^MH^ΔRD and GLI3^R1537C^ΔRD significantly increased reporter activity at comparable levels (Figure 3D). Furthermore, we corroborated that overexpression of GLI3^MH^ΔRD and GLI3^R1537C^ΔRD induced the expression of Olig2, a direct downstream target of GLI proteins, in the developing chick neural tube (Figure 3E). These data indicate that R1537C does not affect the function of transcriptional activator domain of GLI3.

### R1537C affects unique downstream gene expression related to anatomical structures

Our luciferase assay suggest that R1537C affects transcriptional activity of full-length of GLI3. To elucidate the impact of R1537C on GLI3-dependent unique target genes, we overexpressed full-length GLI3^MH^ or GLI3^R1537C^ in HEK293T cells and performed RNA sequencing (RNA-seq) analysis. Because GLI3 is weakly expressed in HEK293T cells, we also performed RNA-seq with control HEK293T cells transfected with empty vector to capture background level of gene expression. First, we checked the expression of direct downstream genes of Hedgehog signaling, such as NKX2.2, NKX6.1, PAX6, IRX3, and OLIG2. Compared to the control, these genes were slightly up- or downregulated by full-length GLI3 overexpression. Importantly, we could not detect differences in Fc of these genes between GLI3^MH^ and GLI3^R1537C^ overexpression (Figure 4A), corroborating that R1537C does not interfere with Hedgehog-dependent signaling. Next, we focused on differentially expressed genes [DEGs; |fold change (Fc)|≥1.5, false discovery rate (FDR)<0.05] between the control and GLI3^MH^ or GLI3^R1537C^ overexpression. Compared to the control HEK293T cells, a total of 846 and 671 genes were differentially expressed by the overexpression of GLI3^MH^ or GLI3^R1537C^, respectively (Table S1). We further conducted Gene Ontology (GO) enrichment analysis to identify the functional characteristics of DEGs. Compared to control, GLI3^MH^ overexpression enriched GO terms related to various biological processes, consistent with the pleiotropic functions of GLI3. Specifically, GLI3^MH^ overexpression enriched GO terms associated with developmental process, such as ossification, renal system development, and extracellular matrix organization (Figure 4B). In contrast, GLI3^R1537C^ did not enriched these GO terms, but increased terms related to chromatin and nucleosome assembly (Figure 4B). To further investigate altered gene regulations by R1537C, we directly compared transcriptomes with GLI3^MH^ and GLI3^R1537C^ overexpression and identified 91 DEGs that were significantly up- or downregulated by GLI3^R1537C^ (DEGs in GLI3^R1537C^ vs. GLI3^MH^; |Fc|>=1.5, FDR<0.05, Table S2). These DEGs included unique transcripts, such as LINC00294, a human specific noncoding RNA that is involved in cell proliferation (33–35), and SLC9A3 (NHE3), a member of the sodium-hydrogen exchanger family that is responsible for the maintenance of sodium balance (36). Among them, 23 genes were associated with developmental process (Figure 4C). Notably, H4 clustered histone 3 (H4C3; Fc=3.7935, p=0.0465) plays an essential role in chromatin organization and disruption of this gene results in growth retardation and skeletal abnormalities in humans (37). Stanniocalcin 1 (STC1; Fc=-2.93183, p=7.15551E-05) is a glycoprotein involved in calcium and phosphate metabolism, and STC1 transgenic mice exhibited dwarf phenotype due to altered osteogenesis (38). These data suggest that the missense mutation of GLI3 alters transcriptional regulations related to specific organ development during embryogenesis.

**Figure 4.**
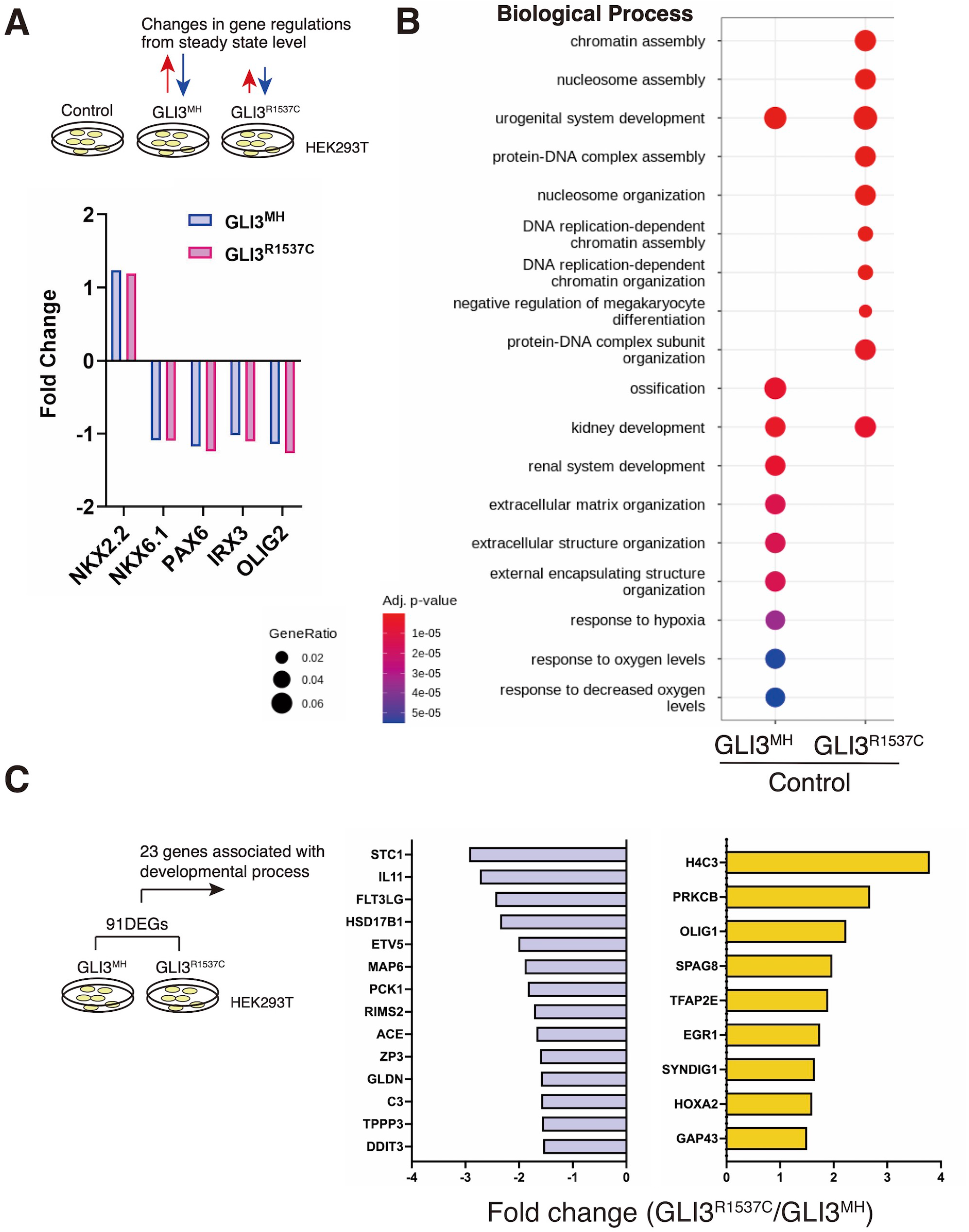
R1537C alters the regulation of GLI3-dependent gene expression. (A) Fold changes of direct downstream genes of Hedgehog signaling in HEK293T cells overexpressing GLI3^MH^ or GLI3^R1537C^ compared with control samples. (B) GO enrichment analysis in comparisons of HEK293T cells overexpressing GLI3^MH^ or GLI3^R1537C^ versus control samples. Spots represent the top 20 ranking terms with gene ratios over 0.02. (C) Genes associated with developmental process that were down- or upregulated by GLI3^R1537C^ compared with GLI3^MH^ overexpression.

### Mice with a Neanderthal/Denisovan-type Gli3 variant exhibit altered skeletal morphology

To further characterize the effect of R1537C on embryonic development, we created knock-in mice with the Neanderthal/Denisovan GLI3 variant by CRISPR-mediated genome engineering technology. In mice, the amino acid that corresponds to the missense variant is naturally arginine (1540R) (Figure 5A, 1A). Thus, we introduced the point mutation c4618t by Cas12 (Cpf1)-dependent homologous recombination, resulting in an R1540C substitution in the mouse GLI3 protein (Figure 5B, S1A-C). Western blotting confirmed that the amino acid substitution did not compromise protein expression compared to wild type GLI3 (1540R) (Figure S1D). On the C57BL6 background, mice carrying 1540C were born at the expected Mendelian ratio. Homozygous 1540C mice had similar body sizes to wild type or heterozygous (1540R/1540C) littermates (Figure S1E). However, we observed altered head morphology in 1540C homozygous mice compared with the wild type or heterozygous littermates. Whole mount skeletal analysis indicated enlarged crania with frontal and parietal bossing in the 1540C mice (Figure 5C, F, Table S3). Previous studies reported that Gli3 null mice (*Gli3^xt/xt^*) exhibited craniosynostosis, a premature closure of cranial sutures, which frequently results in abnormal shape of skulls (39). Consistently, deformed skulls in 1540C mice were frequently associated with premature ossification at the cranial suture (Figure 5D, E). Some homozygous 1540C mice exhibited asymmetric shapes of rib cages associated with scoliosis (Figure 5G, Table S3). Furthermore, compared to wild type mice, the number of lumbar vertebrae was reduced in heterozygous and homozygous mice carrying the 1540C allele (Figure 5H, Table S3). Polydactyly was shown to be a typical phenotype caused by Gli3/GLI3 null conditions in both mice and humans (30). In contrast, the heterozygous and homozygous 1540C mice did not exhibit any abnormalities in limb and digit formation (Figure S1E). These results indicated that a Neanderthal/Denisovan Gli3 variant resulted in altered skeletal morphology, which differed from the phenotypes associated with the loss of Gli3 functions.

**Fig. 5.**
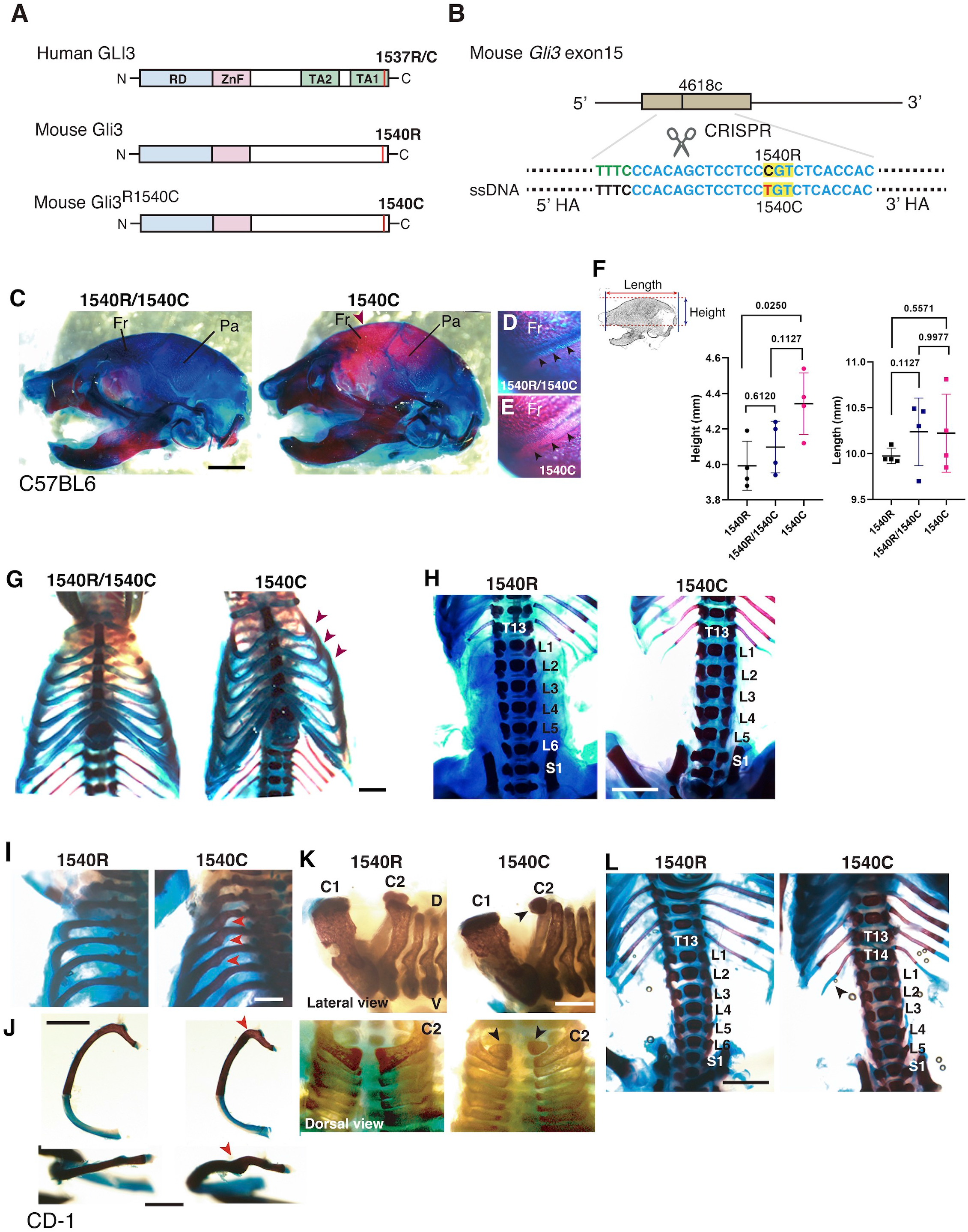
Altered skeletal morphology of the mice carrying a Neanderthal/Denisovan type GLI3 variant. (A) Structures of modern human or extinct hominins GLI3, wild type mouse Gli3 (Gli3^1540R^), and mouse Gli3 with a Neanderthal/Denisovans variant (Gli3^1540C^). (B) Single nucleotide substitution in mouse Gli3 by CRISPR-mediated genome editing. (C-E) The skull of heterozygous (1540R/1540C) and homozygous (1540C) mice on C57BL6 background. An arrowhead indicates an enlarged cranium in 1540C mouse. (D, F) Coronal sutures (arrowheads) of heterozygous (D) and homozygous (E) mice. (F) Quantification of skull height and length. (G) Comparison of rib cages. Arrowheads indicate abnormal rib torsion in 1540C mice. (H) Reduced number of lumber vertebrae in 1540C mouse. (I-L) Skeletal morphology of mice on CD-1 background. The 1540R mice exhibit a twist of the posterior angle of the rib (arrowheads in I and J), hypoplastic ossification in the 2^nd^ cervical vertebrae (arrowheads in K), extra thoracic ribs (T14, arrowheads in L) and reduced number of lumber vertebrae (L). Scale bars: 1 mm.

Several lines of studies on archaic human genomes indicated that Neanderthals and Denisovans had distinct genetic backgrounds compared to modern humans, although these extinct hominins and ancestral modern humans had frequently intermixed with each other (12, 14, 16, 17). Accordingly, the phenotypic consequences caused by the missense variant might be variable depending on specific genetic backgrounds. To examine this possibility, we introduced the 1540C substitution into mice with a CD-1 (ICR) background, which is an outbred strain with heterogeneity of individual genomic sequences compared to inbred strains. Homozygous 1540C mice were viable and their body sizes were similar to those of wild-type or heterozygous knock-in mice (Figure S1F-H). As in the case of the knock-in mice on the C57BL6 background, CD-1 mice carrying the 1540C allele exhibited abnormal rib torsion (Figure 5I, J). In particular, the posterior angle of the ribs in 1540C mice was abnormally kinked, which resulted in the crank-shaft sternum (Figures 5J, S1I and Table S3). Furthermore, we found additional phenotypes in CD-1 background mice, such as hypoplasia of the second cervical vertebrae and extra ribs at the 14^th^ thoracic vertebrae (Figure 5K, L). The number of lumber vertebrae was reduced in the mice with extra ribs (Figure 5L). These skeletal abnormalities were also observed in a lower proportion of the heterozygous mice, indicating a dose-dependency of the phenotypes correlated with the presence of the 1540C allele (Table S3). Curiously, extra 14^th^ ribs occurred in a small proportion of the wild-type CD-1 mice (Table S3), suggesting that the missense Gli3 variant increased phenotypic variations sustained in the background population.

### Gli3^R1540C^ alters the expression of histone cluster genes in mice

To identify downstream genes affected by the missense Gli3 variant in mice, we isolated mesenchymal tissues of wild-type and 1540C mouse embryos on the C57BL6 background and performed RNA-seq analysis. A total of 453 and 238 genes were up- or down-regulated in 1540C compared to wild-type mice [|fold change (Fc)|≥1.5, false discovery rate (FDR)<0.05, Table S4]. We found that DEGs included Msx1, Dlx1 and Dlx2, which were known to be essential for skeletal morphology (40, 41). Notably, several histone H4 cluster genes, such as H4c2, H4c6 and H4c14 were upregulated in 1540C mice, suggesting that the arginine to cysteine substitution of Gli3 commonly affects specific histone components in mice and humans (Table S4).

### Complete disruption of the C-terminal region of Gli3 causes severe abnormalities in developing mouse embryos

To investigate whether the phenotype of the 1540C mice is caused by the disorganization of the C-terminal domain of Gli3, we generated additional mice with a deletion allele of Gli3 on CD-1 background, where a frame shift mutation (R1540Sfs54X) completely disrupts the natural amino acid sequences of the C-terminal region (Figures 6A, S2A-D). Although heterozygous mice (1540Sfs/+) did not show obvious abnormalities, homozygous mice (1540Sfs/1540Sfs) exhibited severe exencephaly and preaxial polydactyly with higher penetrance (Figure 6B-D, Table S3), which were identical to the phenotypes of Gli3 null mutants but differed from 1540C-dependent phenotypes. Together, these results suggest that 1540C substitution increased anatomical variations depending on strain-specific genetic backgrounds, which is attributed to subtle functional alterations of the Gli3 C-terminal domain (Figure 6E).

**Fig. 6.**
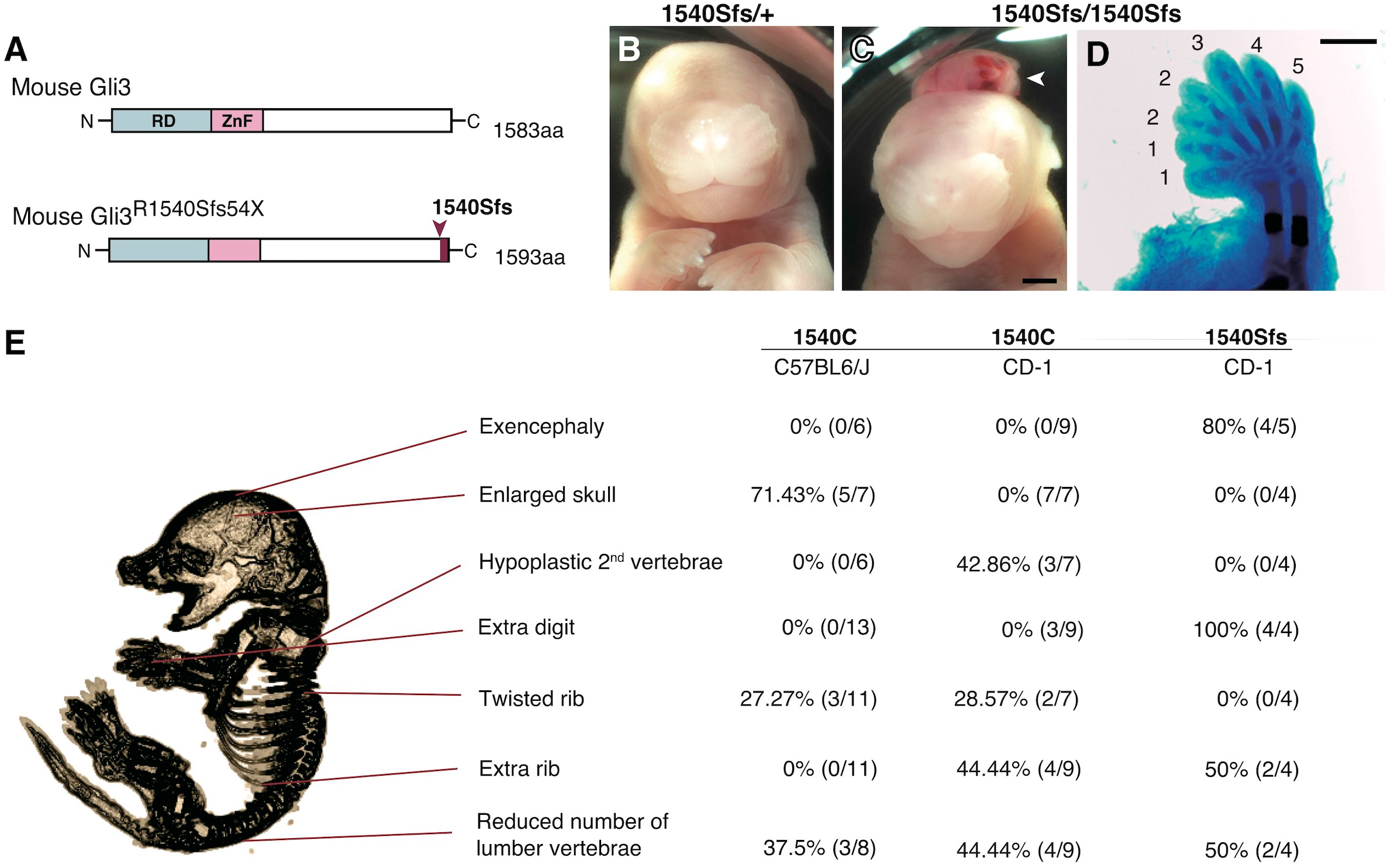
Altered skeletal morphology of Gli3^R1540Sfs54X^ mice and summary of skeletal phenotypes in mice carrying distinct Gli3 variants. (A) A scheme of mouse Gli3^R1540Sfs54X^. (B-D) Phenotypes of heterozygous (1540Sfs/+) and homozygous (1540Sfs/1540Sfs) mice. Homozygous mice exhibit exencephaly (C) and polydactyly (D). (E) Variation of skeletal phenotypes in Gli3^R1540C^ and Gli3^R1540Sfs^ mice. The proportion represents phenotypes in homozygous mice. Scale bars: 1 mm (C), 2 mm (D).

## Discussion

Here, we show that an extinct hominin-specific amino acid alters GLI3 functions by conferring differential transcriptional regulation of downstream genes and results in altered skeletal morphology in mice. Several studies have shown that the ratio of GLI3 activator versus repressor is critical for mediating Hedgehog-dependent signaling during embryogenesis (42, 43). Despite *in silico* prediction, we demonstrate that R1537C substitution does not compromise GLI3 protein stability or activator-dependent transcriptional activity. Furthermore, R1537C does not alter the regulations of direct target genes of Sonic Hedgehog signaling. Thus, GLI3^R1537C^ does not interfere with the potential to mediate Hedgehog signaling during embryogenesis, thereby detouring deleterious phenotypes. In contrast, disruption of the Gli3 C-terminal region causes severe exencephaly and polydactyly in mice, as in the case of the pathogenic mutations of human GLI3. Therefore, the presence of the C-terminal region is a prerequisite for GLI3 functions, possibly through interaction with various cofactors for transcriptional regulation (44).

We demonstrate that mice with a Neanderthal/Denisovan GLI3 variant exhibit altered skeletal structures, such as enlarged cranium, altered shapes of vertebrae, and rib malformations. A recent study reported the association of unique anatomical phenotypes in Neanderthal with Human Phenotypic Ontology (HPO), a database of disease-associated human phenotypes (45). Notably, some of neanderthal-specific HPO phenotypes, such as macrocephaly (HP: 0000256), accelerated skeletal maturation (HP: 0005616) and scoliosis (HP: 0002650) are linked with the phenotypes of R1540C mice (Table S5). We find that the mice carrying the Neanderthal/Denisovan GLI3 variant also showed altered shapes of rib cages due to abnormal rib torsion. Fossil evidence indicated a stronger torsion of central ribs (6^th^-8^th^ ribs) in Neanderthal infants, which contributes to distinct rib cages (shorter and deeper forms) compared to those of modern humans (46). Even if not all skeletal phenotypes of R1540C mice phenocopied anatomical characteristics of extinct humans, phenotypic alterations in the knock-in mice suggest that the corresponding missense mutation of GLI3 might affect conserved developmental processes of homologous skeletal structures between mice and humans.

The corresponding amino acid substitution is highly unique in archaic hominins and modern humans. Various skeletal abnormalities induced by archaic hominin-type substitution might not be adaptive in mice. These lines of evidence suggest that concurrent changes in developmental programs occurring in archaic hominins accommodated R1537C-dependent phenotypic abrogation. Alternatively, the variant-dependent functional alterations might be accepted by relaxed developmental constraint occurred in archaic and modern human lineages. R1537C-dependent phenotypes could be attributed to the interaction with other variants reside in archaic and modern human-specific genetic backgrounds. We confirmed that differential genetic background significantly affects GLI3 variant-dependent phenotypic variations in mice. Phenotypic differences might be due to genetic and/or epigenetic differences in other genomic regions, which affect GLI3-dependent developmental processes. Notably, the R1540C variant frequently contributed to 14^th^ rib formation in CD-1 background, which also occurs occasionally in wild type population. It has been reported that variable number of vertebrae depends on genetic backgrounds as well as maternal influences (47, 48). A recent study reported that the R1537C variant is associated with neural tube defects in European population (49). Thus, it is possible that the GLI3^R1537C^ increased susceptibility to variable morphological characteristics or pathogenic phenotypes in response to genetic and environmental perturbations.

It remains still unclear that R1537C variant was fixed in extinct hominin groups by positive selection. Recent studies have suggested that the small population size of Neanderthals was not effective for natural selection, by which weakly deleterious mutations might be accumulated by genetic drift (12, 13). Notably, datasets of associations between various human traits and variants based on UK Biobank cohorts (http://geneatlas.roslin.ed.ac.uk, https://pheweb.org/UKB-SAIGE/) revealed that R1537C (A allele) is associated with specific anatomical traits, such as anisotropy of the superior longitudinal fasciculus (p=0.00038) waist-hip ratio, (p=0.00057), and other unspecified back disorders (p=0.0000045), while the modern human variant (G allele) increases the risk of lethargy (p=0.00058) and osteoarthritis (p=0.00067). Although most of these associations did not pass statistical correction for multiple comparisons, these traits are linked with the predicted lifestyles of Neanderthals, suggesting that R1537C provided beneficial traits for extinct hominins. To distinguish these possibilities, further analyses are required to identify the biological effects of the variant in human development, as well as genetic and epigenetic modifiers that affect variant-dependent phenotypes.

## Materials and Methods

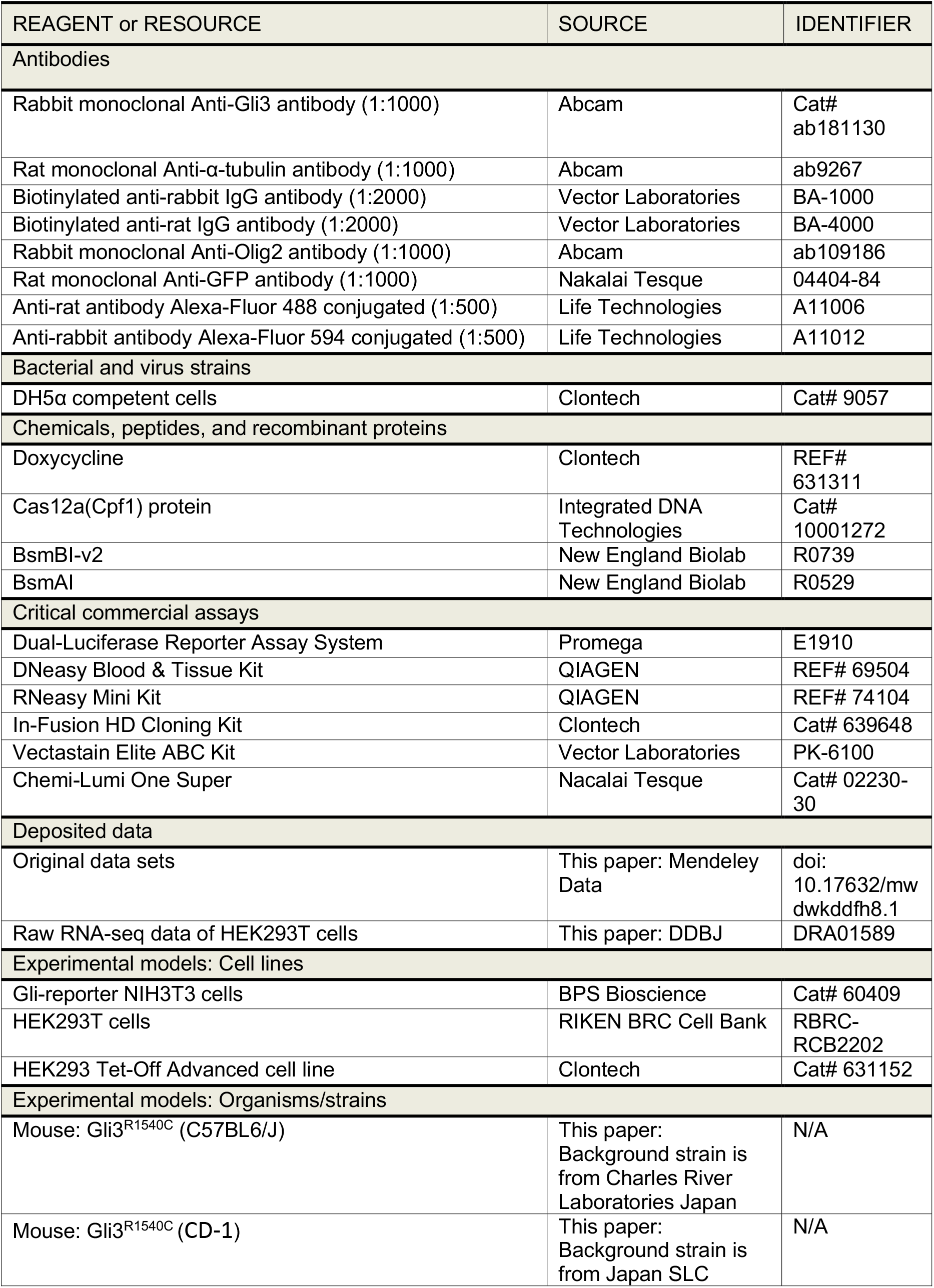
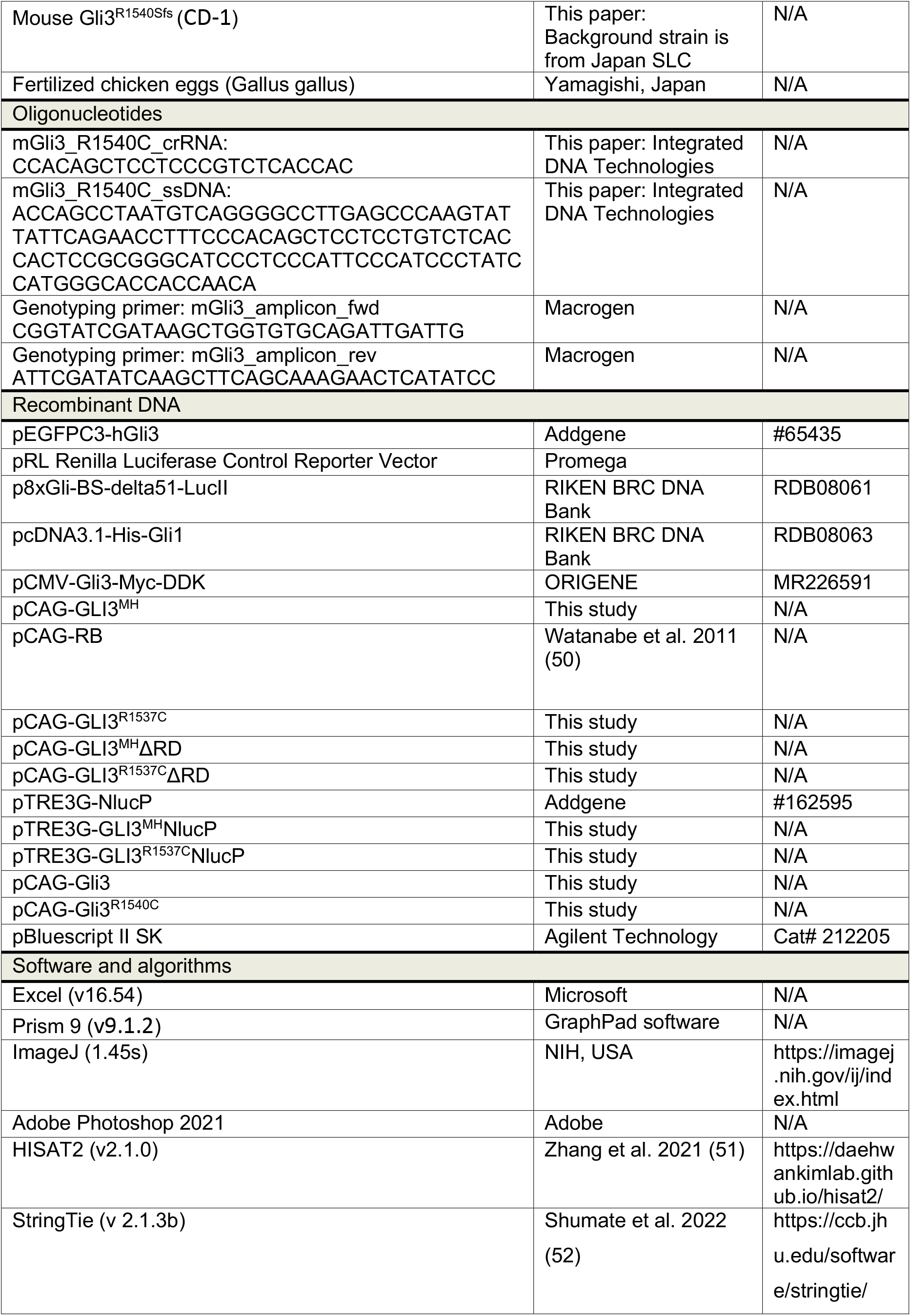
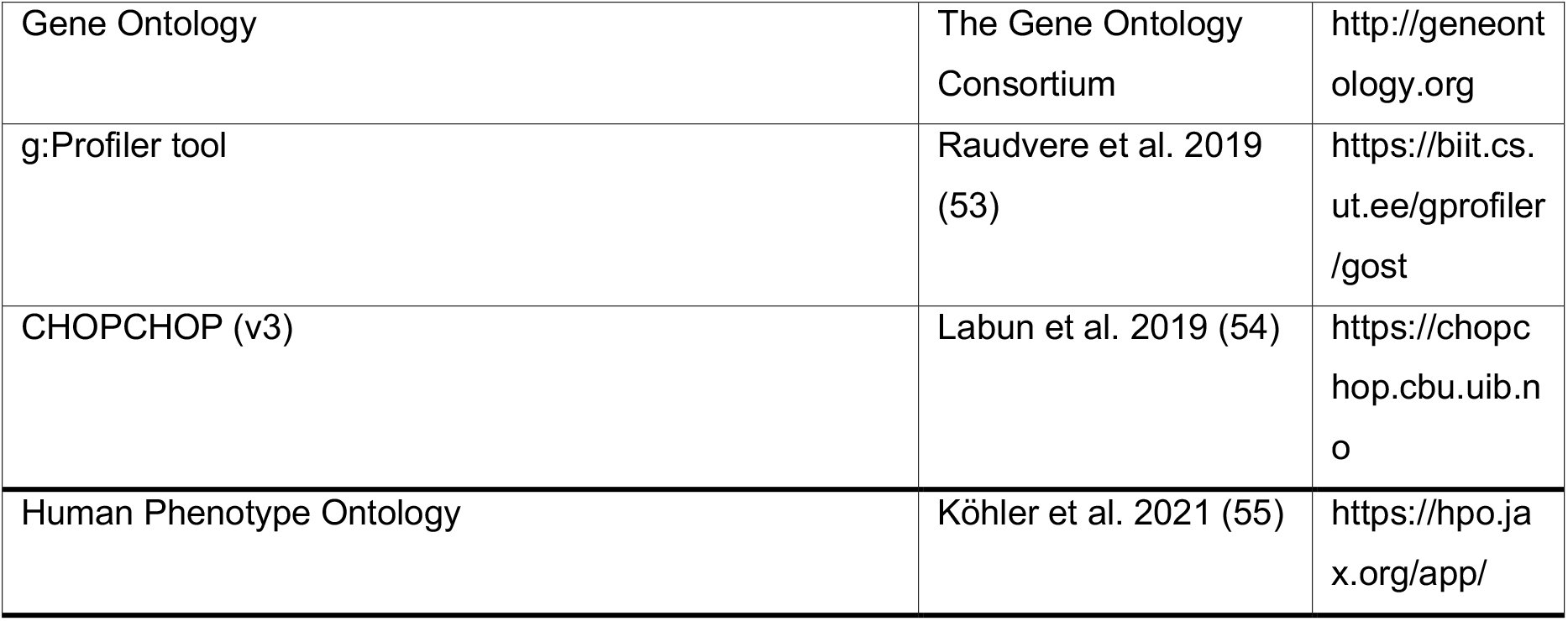

### Animals

Male and female mice (CD-1 and C57BL6/J backgrounds) that were originally obtained from Japan SLC and Charles River Laboratories Japan were maintained in a 12 h dark/light cycle at the Experimental Animal Facility of Kyoto Prefectural University of Medicine. Noon of the day the vaginal plug was identified was designated as E0.5. Fertilized chicken eggs (*Gallus gallus*) were obtained from a local farm (Yamagishi, Japan) and incubated at 37.0 ± 0.2 °C. Embryonic stages were determined according to the Hamburger-Hamilton stages (56). All animal experiments were approved by the experimental animal committee of Kyoto Prefectural University of Medicine and were performed in accordance with the relevant guidelines (M2021–233, M2022-209, M2021-511, M2022-210, M2021-217, M2022-201).

### Plasmids

Full-length human GLI3 and mouse Gli3 cDNAs were obtained from pEGFPC3-hGli3 (a gift from Aimin Liu, Addgene plasmid, (57)) and pCMV-Gli3-Myc-DDK (ORIGENE, MR226591), and then subcloned into the pCAG-RB vector by using an In-Fusion HD cloning kit. For construction of pCAG-GLI3^R1537C^ and pCAG-mGli3^R1540C^, the 3’ regions of GLI3 or mGli3 were replaced with PCR-amplified fragments containing c4609t (GLI3^R1537C^) or c4618t (mGli3^R1540C^), respectively. pCAG-GLI3^MH^ΔRD and pCAG-GLI3^R1537C^ΔRD were generated by subcloning the C-terminal region of GLI3^MH^ or GLI3^R1537C^ into the pCAGRB vector, according to the method of a previous report (58). For construction of pTRE-GLI3^MH^-NLucP and pTRE-GLI3^R1537C^-NLucP, coding sequences of GLI3^MH^ or GLI3^R1537C^ were subcloned into pTRE3G-NlucP (a gift from Masaharu Somiya, addgene plasmid, (59)). All vectors were verified by sequencing.

### Luciferase reporter assay

Gli-reporter NIH3T3 cells were transfected with pCAG-GLI3^MH^ or pCAG-GLI3^R1537C^ together with pTK-RL by using Lipofectamine 2000 (Thermo Fisher Scientific). To monitor Gli-dependent luciferase activity in HEK293T cells, p8xGli-BS-delta51-LucII (60) was transfected with other expression vectors. As a control condition, the pCAGRB empty vector was cotransfected with reporter vectors. For analysis of GLI1-dependent reporter activities, pcDNA3.1-His-Gli1 (61) was transfected with GLI3 expression vectors. Transfected cells were cultured in Dulbecco’s modified Eagle’s medium (high glucose, Nacalai Tesque) containing 10% fetal bovine serum and 1% penicillin and streptomycin (Fujifilm Wako Pure Chemical Corporation) for 24 h. For analysis of the degradation rates of GLI3^MH^ and GLI3^R1537C^, pTRE-GLI3^MH^-NLucP or pTRE-GLI3^R1537C^-NLucP was transfected into the HEK293 Tet-Off Advanced cell line, and doxycycline was then added to the culture medium 12 hours after transfection. Luciferase activity was examined with the Dual-Luciferase Reporter Assay System and analyzed with a luminometer (GENE LIGHT GL210A, Microtec, Inc.). All firefly luciferase values were normalized to Renilla luciferase activities. At least three biologically independent samples were analyzed in each experimental condition, and all experiments were confirmed by at least three technical replicates.

### Western blotting

HEK293T cells transfected with pCAG-GLI3^MH^, pCAG-GLI3^R1537C^, pCAG-mGli3, or pCAG-mGli3^R1540C^ were lysed in RIPA buffer (20 mM Tris, 150 mM NaCl, 1 mM EDTA, 1% Nonidet P40, 1% SDS, 0.1% deoxycholate, 1 mM NaF and protein inhibitor). After electrophoresis, proteins were transferred to Poly Vinylidene Di-Fluoride (PVDF) membranes, blocked with Bullet Blocking One for Western blotting (Nacalai Tesque), and then incubated with anti-human GLI3 or anti-α-tubulin antibody. The membranes were further incubated with biotinylated anti-rabbit IgG or biotinylated anti-rat IgG antibody, followed by a Vectastain Elite ABC kit and developed with Chemi-Lumi One Super and analyzed with a luminescent image analyzer (LAS-2000, Fujifilm).

### *In ovo* electroporation

*In ovo* electroporation of developing chick embryos was performed according to a previous study (58). Briefly, a window was opened in the shell of an egg, and then a small amount of DNA solution (less than 0.05 µL) was injected into the neural tube of E2 (HH stage 13-15) embryos with a fine glass needle. Needle-type electrodes (CUY200S, BEX) were placed on the neural tube and square electric pulses (28 V for 50 ms, 2 or 3 times) were applied to the thoracic neural tube with a pulse generator (CUY21 EDITII, BEX). After electroporation, the extraembryonic cavity was filled with sterilized Hank’s balanced salt solution (HBSS, Nacalai Tesque) containing antibiotics (penicillin/streptomycin and gentamycin), and the window was sealed with parafilm. Electroporated embryos were incubated in an incubator at 37 °C for 24 to 48 hours.

### Immunohistochemistry

Electroporated embryos were fixed with 4% paraformaldehyde dissolved in phosphate-buffered saline (PBS) at 4 °C overnight. After PBS washes, the embryos were cryoprotected with a 20% sucrose solution and immersed in Tissue-Tek. The frozen samples were sectioned at a thickness of 20 µm using a cryostat (Leica CM1850, Germany), and incubated with primary antibodies, including anti-Olig2 and anti-GFP antibodies. After washing, the sections were incubated with secondary antibodies, including Alexa-Fluor 488 or 594 conjugated anti-rat or anti-rabbit antibodies. Fluorescence images were captured with fluorescence microscopes (BX51, Olympus) equipped with a cooled CCD camera (DP80, Olympus). All captured images were processed with cell Sense standard (v.1.17, Olympus), ImageJ and Adobe Photoshop.

### RNA-seq analyses

Total RNA was prepared from HEK293T cells transfected with pCAG-RB, pCAG-GLI3^MH^ or pCAG-GLI3^R1537C^ by using an RNeasy kit. To isolate total RNA from mouse tissue, thoracic mesenchymal tissue of E17.5 wild-type and 1540C mice was dissected and preserved in RNAlater until RNA extraction. Residual DNA was eliminated by DNase treatment. The quality and quantity of RNAs were assessed by using an Agilent Technologies 2100 Bioanalyzer or 2200 TapeStation (Agilent). The cDNA library was constructed using a TrueSeq Standard mRNA LT Sample Prep Kit according to the manufacturer’s protocol (15031047 Rev. E) and sequenced on an Illumina platform and 101 bp paired-end reads were generated. The sequence data were mapped to a reference genome sequence (*Homo sapiens* GRCh38; NCBI_109.20200522 or *Mus musculus* GRCm38, GCA_000001635.2) with a splice-aware aligner (HISAT2 v 2.1.0). The transcripts were assembled by StringTie (v 2.1.3b) with aligned reads. GO enrichment analysis was performed based on Gene Ontology. A significant gene list was constructed by the g:Profiler tool.

### SNP population genetics

The genetic population of rs35364414 (GLI3 4609G/A) was analyzed by Ensemble genome browser 108 (https://www.ensembl.org/index.html).

### Mouse genome engineering

For generation of a point mutation in mouse Gli3 that corresponds to a Neanderthal GLI3 variant, genome edited mouse lines were established by CRISPR/Cas12a(Cpf1)-mediated homology directed repair (HDR). A crRNA sequence that targets mouse Gli3 4618c was designed by CHOPCHOP. We chose a target sequence with no off targets based on mouse genome information (mm10/GRCm38). As an HDR template for homologous recombination, single strand DNA (ssDNA) that carries a nucleotide substation (4618t) was designed by using the mouse Gli3 sequence.

### Generation of genome edited mice

The cumulus complexes were isolated from superovulated female C57BL6/J mice treated with pregnant mare serum gonadotropin (PMSG) and human chronic gonadotropin (hCG), and then mixed with sperm taken from the epididymis of male C57BL6/J mice. After in vitro fertilization, the CRISPR/Cas12a solution containing Cpf1 protein (Integrated DNA Technology, Inc.), crRNA, and ssDNA were introduced into zygotes by electroporation (Genome Editor Plus. BEX, Tokyo, Japan). After electroporation, the zygotes were transferred into the oviduct of pseudopregnant CD-1 mice. Anesthesia was performed by intraperitoneal injection of medetomidine, midazolam and butorphanol. Genome-edited mice on the CD-1 background were generated by the *i*-GONAD method (62). Briefly, pregnant female mice (E0.75) were anesthetized with 2% isoflurane and the CRISPR/Cas12a solution was injected into the oviduct lumen. Then, square electric pulses were applied to the oviduct by using a pulse generator (super electroporator NEPA21, Nepa gene).

### Genotyping of mice

Genomic DNA from embryonic or adult mouse tissue (skin or tail fragments) was prepared using a DNeasy Blood & Tissue kit. The DNA fragments containing the CRISPR/Cas12a target region were amplified by polymerase chain reaction (PCR). PCR fragments were digested by BsmAI or BsmAI for 1 hour, and restriction fragment length was determined by electrophoresis with 3% agarose. For determination of the edited sequences of candidate mouse lines, PCR amplicons were subcloned into the pBluescriptSK vector, and six to ten randomly selected clones were examined by Sanger sequencing. F0 mice heterozygous for the 4618c>t substitution (Gli3^R1537C^) or nucleotide deletion (Gli3^R1540Sfs^) were propagated and used to generate homozygous mice.

### Skeletal analyses

Embryonic and postnatal mice were collected (embryonic Day 16.5, postnatal Day 1, or Day 7) and dissected in phosphate buffered saline. The samples were fixed with 100% ethanol for 3 days and degreased by acetone overnight. Skeletal staining was performed by Alcian blue and Alizarin red according to a standard protocol (63). Sample images were captured by using the microscope (SZX7, Olympus) equipped with a cooled CCD camera (DP80, Olympus).

### Image processing and analyses

Images of Western blotting were examined by using ImageJ. All captured images were processed with Adobe Photoshop 2021. Original Western blotting images were deposited in Mendeley Data.

### Statistical analyses

For statistical analysis, at least three independent samples from each experimental group were compared. Comparisons between experimental groups were performed using Microsoft Excel and Prism 9. All data are presented as the mean ± SE. Statistical significance was determined using the two-tailed unpaired Student’s *t* test, ordinary one-way ANOVA with Tukey’s multiple comparisons test, or two-way ANOVA with Sidak’s multiple comparison test. Statistical analysis of differential gene expression conducted by RNAseq data was performed using fold change and the nbinomWald Test using the DESeq2 R package (Bioconductor) per comparison pair, and significant results were selected based on |fc|≥1.5 and nbinomWald Test with a raw p value <0.05.

## Author contributions

A.A. and T.N. contributed to the experimental design, data collection, interpretation, and manuscript preparation. S.O. contributed to the generation and maintenance of mice carrying GLI3^R1540C^. R.N. contributed to the collection and analysis of RNA-seq samples, H.G. and K.O. contributed to the acquisition of the reagents and interpretation. All authors discussed the data and interpretation and confirmed the final version of the manuscript.

## Acknowledgements

We thank Drs. Ikuo K. Suzuki and Takuma Kumamoto for critically reading the manuscript and Ms Kawami Misato and Ms Mariko Yazaki for technical assistance.

## Data availability

The raw RNA sequencing data have been deposited in the DDBJ database (DRA015189 and DRA015774). All data generated in this study have been deposited in Mendeley Data (DOI: 10.17632/mwdwkddfh8.2).

## Supplementary Items

Table S1. DEGs in GLI3^MH^ or GLI3^R1537C^ vs. control HEK cells

Table S2. DEGs in GLI3^MH^ vs. GLI3^R1537C^ HEK cells

Table S3. Skeletal phenotypes of mice carrying GLI3^R1540C^

Table S4. DEGs in wild-type vs. GLI3^R1540C^ mice

Table S5. Comparisons of HPO phenotypes associated with Neanderthals and mice carrying GLI3^R1540C^.

